# NITINOL MATERIAL PROPERTIES OF 11 COMMERCIAL PERIPHERAL STENTS DETERMINED USING INVERSE COMPUTATIONAL ANALYSIS

**DOI:** 10.1101/2025.11.27.691034

**Authors:** Eric Anttila, Kaspars Maleckis, Majid Jadidi, Anastasia Desyatova, Jason MacTaggart, Alexey Kamenskiy

## Abstract

Stent-artery interactions are influenced by the mechanical properties of self-expanding Nitinol stents, but data on these characteristics remain limited. Eleven stents (Absolute Pro, S.M.A.R.T. Control, Misago, Zilver, Complete SE, EverFlex, Innova, Pulsar-18, LifeStent, S.M.A.R.T. Flex, and Supera) used to treat peripheral arterial disease (PAD) were subjected to axial tension, compression, three-point bending, and torsion tests, and the data on reaction forces and moments were compared with finite element simulations of the same experiments. Inverse computational analysis was used to determine austenite and martensite elasticity, transformation stretch, stresses at the start and end of transformation loading, and the start of transformation stress in compression. Uniaxial tensile tests were done on isolated struts from Absolute Pro and Zilver stents to verify the results of the inverse analysis. Our study demonstrate that Nitinol material properties are significantly different across devices. Austenite elasticity ranged 7.5-85 GPa, martensite elasticity 10-47.8 GPa, transformation stretch 1.03-1.08, the start of transformation loading stress 386-465 MPa, the end of transformation loading stress 411-535 MPa, and the start of transformation stress in compression 150-900 MPa. Nitinol of S.M.A.R.T. Control and S.M.A.R.T. Flex devices had the softest response, while Pulsar-18 had the hardest. The presented Nitinol mechanical properties of commonly used PAD stents can improve the fidelity of computational models investigating stent-artery interactions and may help improve clinical outcomes of endovascular PAD repairs through better device design.

## 1. INTRODUCTION

Peripheral Arterial Disease (PAD) commonly refers to the flow-obstructing atherosclerotic narrowing of the femoropopliteal artery (FPA) in the lower limb^1^. It is associated with high morbidity, mortality, and significant quality of life impairment^2–4^. PAD patients are at significantly higher risk for major cardiovascular events and are four times more likely to die within a 10-year time period than similarly-aged non-PAD subjects^5^. PAD healthcare expenditures exceeded $21 billion in the United States alone^6^, and a significant part of these costs is attributed to the high number of repeat interventions to treat in-stent restenosis^7–11^. Though the exact reason for poor clinical results that require these interventions is not completely understood^12^, the interactions between the repair device and the dynamic environment of the FPA that experiences complex deformations with limb flexion are thought to play an important role^13–15^.

Balloon angioplasty, often followed by stenting, is one of the most common minimally invasive PAD treatments. During angioplasty, a balloon catheter is inserted via a small incision in the groin and guided to the obstructed segment of the artery under fluoroscopy. The balloon is then inflated, crushing the plaque and restoring blood flow to the distal tissues. While in many cases balloon angioplasty is sufficient to resolve the occlusion, flow-limiting arterial dissections resulting from the delamination of the plaque are not uncommon^16–18^, and require a stent to push the dissection septum out of the way of the flow^19,20^. PAD stents are metal mesh tubes that act as a scaffold to keep the arterial lumen open. They are made of Nitinol, which is a shape memory alloy that combines nickel and titanium to achieve superelastic properties. Nitinol stents are self-expanding, biocompatible, and are able to undergo large elastic deformations returning to their pre-set shape. These characteristics make Nitinol stents particularly suitable for the lower extremity applications^21^ as the stented artery undergoes large deformations during flexion of the limbs^22^.

The unique shape memory and superelastic properties of Nitinol stem from the changes in its crystalline structure that can undergo phase transitions under mechanical or thermal loads. Multiple factors can influence the material properties of Nitinol, including heat setting temperature, time, surface treatments, and cooling rate^21^. Stent manufacturers manipulate these characteristics to achieve the desired behavior for their devices, but the specific mechanical properties of their Nitinol remain largely proprietary. This complicates computational assessments of stent-artery interactions that often have to rely on the “generic” Nitinol characteristics^23–34^. Few studies attempted to investigate the differences between Nitinols used in commercial PAD stents^30,33,35^, but the mechanical experiments on individual stent components are challenging because these devices rarely contain large straight segments suitable for reliable mechanical testing.

In the current study, we have determined Nitinol material properties of 11 commercial PAD stents using a combination of inverse computational analysis and experimental benchtop evaluations of the entire devices under axial tension, compression, torsion, and three-point bending deformations. These results were then validated with the single-strut stent experiments for the two commercial devices that had suitable geometry. Our data illustrate the differences between Nitinols used in commercial PAD stents, and the summary of their mechanical properties can be used to improve the fidelity of computational models investigating stent-artery interactions. These models may help improve PAD clinical outcomes through better device design.

## 2. MATERIALS AND METHODS

### 2.1 Reverse engineering of stent geometries

A total of 11 commercially available Nitinol PAD stents were selected for analysis. These included Absolute Pro (Abbott Vascular, Abbott Park, IL, USA), S.M.A.R.T. Control (Cordis, Hialeah, FL, USA), Misago (Terumo, Shibuya City, Tokyo, Japan), Zilver (Cook Medical, Bloomington, IN, USA), Complete SE (Medtronic, Dublin, Ireland), EverFlex (Medtronic, Dublin, Ireland), Innova (Boston Scientific, Marlborough, MA, USA), Pulsar-18 (Biotronik, Berlin, Germany), LifeStent (Bard, New Providence, NJ, USA), S.M.A.R.T. Flex (Cordis, Hialeah, FL, USA), and Supera (Abbott Vascular, Abbott Park, IL, USA). Computer-Aided Design (CAD) models for each device were constructed by rolling the stent on Play-Doh (Hasboro, Rhode Island, USA), optically imaging the imprint left by the stent, and digitizing it in SolidWorks (Dassault Systemes Corp., Rhode Island, USA).

### 2.2 Stent mechanical evaluation

All stents were mechanically tested under axial tension/compression, torsion, and three-point bending at 37°C under displacement-controlled cyclic loading, and the last cycle and the highest peak deformation were used for analysis^36^. Axial tension and compression tests were performed with a CellScale BioTester (Waterloo, Ontario, Canada) and stents mounted on cylindrical supports and clamped around the perimeter with a plastic barb. Span lengths between the clamps varied from 19.5 mm to 27.0 mm. Tests were completed with a 0.467 mm/s displacement rate to 25% tension and compression relative to the span length between the clamps for each stent. Torsion tests were performed in a temperature-controlled air chamber using a TA Electroforce 5175 BioDynamic tester (New Castle, DE, USA). Stents were mounted using tapered supports and clamped with a non-adhesive Parafilm tape, thus restricting all movement at the ends of the stent. The average span between the clamps was 23.0 ± 1.4 mm to comply with previous measurements^14,36^. Cyclic twists of 45°/cm and 90°/cm were applied at 5°/s rotational speed. Three-point bending tests were performed in a horizontal plane using a CellScale BioTester. Stents were supported vertically by a baseplate and horizontally by two 6 mm stainless steel metal pillars located 34 mm apart from one another. A 10 mm high and 6.35 mm wide rounded tip was used to drive the loading pin and simulate bending at the middle segment of the stent. The displacement rate was set to 1.0 mm/s.

### 2.3 Model information

CAD stent geometries were imported into Abaqus CAE (Dassault Systemes, Simulia Corp., Rhode Island, USA), and the analysis was conducted using Abaqus/Explicit. In order to recreate our benchtop tests, the clamps were simulated as rigid bodies with datum planes separating the clamped region from the testing region at equal distances informed by the experiments (range 19.5 to 27.0 mm). A semi-automatic stable time increment ranging from 1e-07 to 1e-09 was used, depending on the stent geometry and mesh. This increment ensured the ratio of kinetic to internal energy did not exceed 5%, thus avoiding dynamic effects in the simulations. All stents were meshed with 8-node linear brick reduced integration elements (C3D8R), and the number of elements for each device was determined with a sensitivity analysis to ensure that the reaction force changed less than 5%.

In order to simulate tension/compression and torsion, boundary conditions were applied via the reference point of the rigid body of one of the clamps, while the other clamp was restricted from all translations and rotations. For axial tension and compression, displacement values were calculated as ±25% of the middle region and applied to the reference point. For torsion, a boundary condition of 90°/cm was calculated based on the length of the middle region and applied to the reference point. For three-point bending, the loading pin was displaced 10 mm to replicate the experimental tests and achieve a 10 mm bending radius. For axial tension/compression and torsion, self-contact, a penalty method, and a frictional coefficient of 0.3 were used. For three-point bending, contact was defined between the stent, the two metal pillars, and the loading pin to replicate experimental conditions. Likewise, a penalty method was used with a frictional coefficient of 0.3. Reaction forces and reaction moments were measured at the clamped end that was restricted from all translations and rotations. These values were then compared with the experiment.

### 2.4 Nitinol parameter determination using the inverse method

After simulating each of the four mechanical deformations, the Nitinol material properties were adjusted until one set of parameters matched reaction forces and moments across all deformations. For this, stents were first assigned a generic set of commonly used Nitinol material properties^32^, and the devices were subjected to axial tension. FEA results were then compared to the experimental axial tension reaction force, and the Nitinol material properties were adjusted until calculations matched the experiments. These material properties included austenite elasticity (*E*_*A*_), martensite elasticity (*E*_*M*_), transformation stretch (*λ*^*L*^), the start 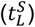 and end 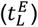 of transformation loading stress, and the start of transformation stress in compression 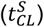. Note that the start and end of transformation stresses in the unloading were not determined in this study, as all stents were only loaded to their final deformed configurations. Next, axial compression followed by torsion and three-point bending tests were simulated with the newly determined material properties, and the parameters were once again adjusted in an iterative fashion. This process continued until reaction forces and moments determined by FEA matched the experimental results with a combined average error of less than 15% in all four deformation modes.

### 2.5 Uniaxial extension of straight Nitinol bridge connectors from Absolute Pro and Zilver stents

Abbott Absolute Pro and Cook Zilver stents had struts sufficiently long and straight to separate and evaluate experimentally. Straight bridge connector struts from these stents were excised, and the ends of the struts were embedded into hard epoxy. The resulting structure was mounted using metal clamps onto the CellScale BioTester equipped with 10 N loadcells. The samples were submerged into a 37°C water bath and subjected to uniaxial extension at 0.01 s^-1^ strain rate up to the 10 N maximum limit to capture the martensitic phase. In-plane stretch was measured in the longitudinal direction (*λ*_*z*_) and related to the corresponding load (*P*_*z*_). The through-thickness average experimental Cauchy stress (kPa) was then calculated as:

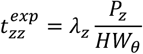

where *H* is the undeformed thickness of the strut measured using a caliper prior to the experiment, and *W*_*θ*_ is the undeformed specimen width over which the respective load *P*_*z*_ acts. The experimental stress-stretch data were then imported into Abaqus CAE, and its calibration module was used to determine the Nitinol material properties presented in Table 1 and schematically illustrated in Figure 1. These material characteristics were then used to validate the Nitinol material parameters determined for the Absolute Pro and the Zilver stents using the inverse approach.

**Table 1.** List of Nitinol material characteristics. Reference temperature was 37°C.

| **Determined Nitinol material characteristics** |
| --- |
| Austenite elasticity, $E_A$ (GPa) |
| Martensite elasticity, $E_M$ (GPa) |
| Transformation stretch and volumetric transformation stretch, $\lambda^L$ |
| Start of transformation loading, $t_L^S$ (MPa) |
| End of transformation loading, $t_L^E$ (MPa) |
| Start of transformation stress in compression $t_{CL}^S$ (MPa) |
| **Other Nitinol characteristics:** |
| Austenite Poisson's ratio, $\nu_A = 0.33$ |
| Martensite Poisson's ratio, $\nu_M = 0.33$ |
| Loading, $(\delta t / \delta T)_L = 0$ (MPa T<sup>-1</sup>) |
| Unloading, $(\delta t / \delta T)_U = 0$ (MPa T<sup>-1</sup>) |
| Start of transformation unloading, $t_U^S = 227$ (MPa) |
| End of transformation unloading, $t_U^E = 187$ (MPa) |

**Figure 1.**
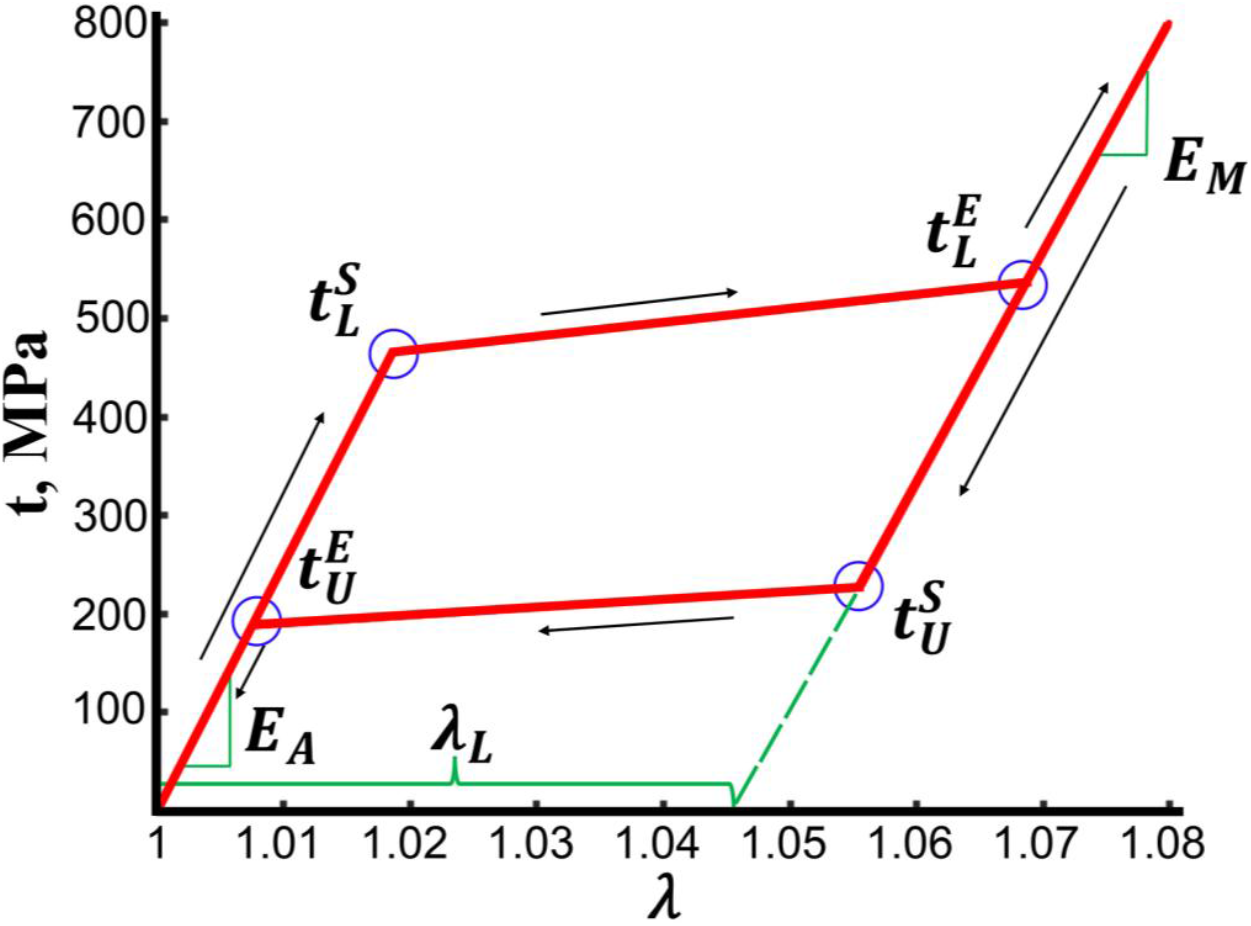
Schematic of the superelastic Nitinol stress-stretch response during loading and unloading (red) and the material characteristics that determine this behavior.

## 3. RESULTS

Reverse-engineered stent geometries depicted next to the photos of the actual devices are demonstrated in Figure 2.

**Figure 2.**
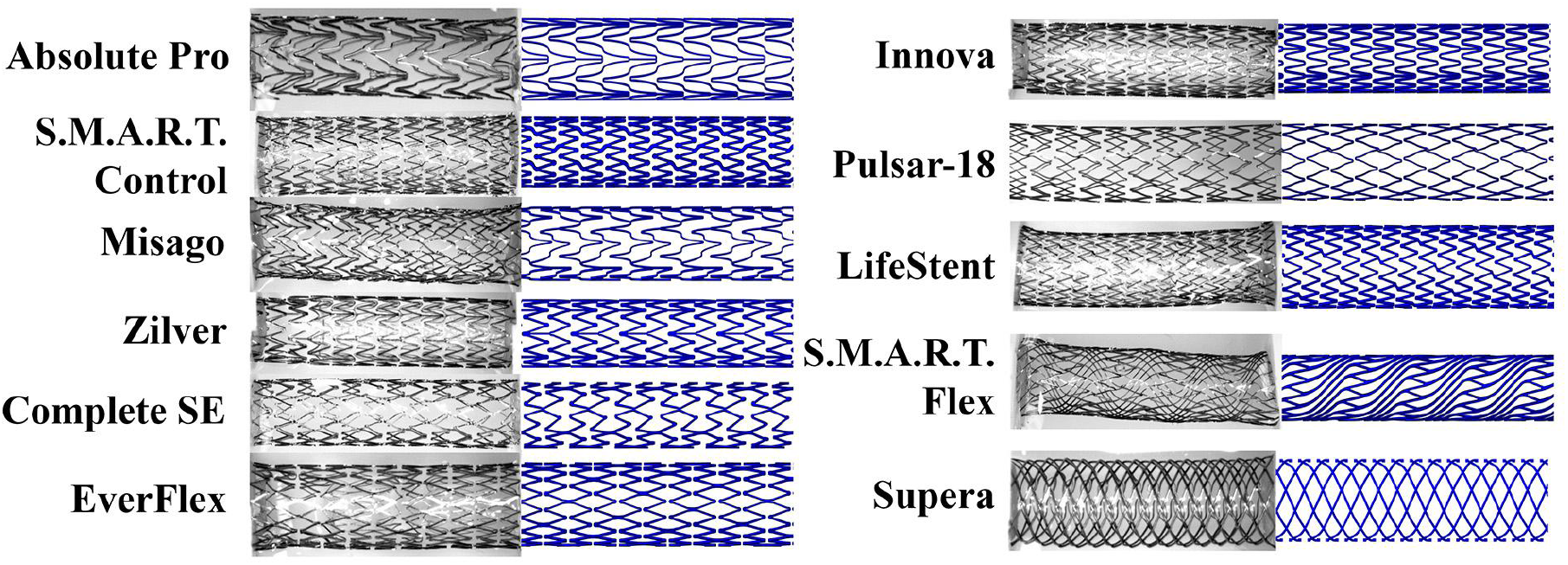
Reverse-engineered stent geometries (right) plotted next to the photos of the actual devices (left).

Results of the 25% axial tension and compression test are presented in Figure 3 and Figure 4, and the experimental and FEA-predicted values of reaction forces are summarized in Table 2. The largest reaction forces under axial tension were observed for the S.M.A.R.T. Flex (Exp 8.70 N vs FEA 9.89 N) and S.M.A.R.T. Control (Exp 2.60 N vs FEA 2.74 N) stents, while the smallest were for the Supera (Exp 0.28 N vs FEA 0.23 N) and Misago (Exp 0.22 N vs FEA 0.19 N) devices. All stents ranked the same experimentally and computationally, with the exception of Innova that demonstrated a stiffer response than the EverFlex in the FEA, but not under the benchtop tension test. On average, the relative error between the FEA and the axial tension experiment was 10.9±4.3%, and the largest discrepancy was observed for the Supera stent (18%), although the absolute error was only 0.05 N (Table 2).

**Table 2.** Experimental and FEA results showing reaction forces, moments, and the corresponding errors for all 11 stents. Misago, Pulsar-18, LifeStent, and S.M.A.R.T. Flex devices had non-symmetric patterns, and their torsion is reported for the clockwise and counter-clockwise twists separately (CW/CCW).

| Device | Tension (N) |  |  | Compression (N) |  |  | Torsion (N·mm) |  |  | 3-Point Bending (N) |  |  |
| --- | --- | --- | --- | --- | --- | --- | --- | --- | --- | --- | --- | --- |
|  | Exp. | FEA | % error | Exp. | FEA | % error | Exp. | FEA | % error | Exp. | FEA | % error |
| **Absolute Pro** | 0.52 | 0.49 | 5.7 | 0.56 | 0.52 | 7.7 | 0.54 | 0.48 | 11 | 0.15 | 0.14 | 7.1 |
| **S.M.A.R.T. Control** | 2.60 | 2.74 | 5.3 | 2.02 | 2.18 | 7.9 | 4.54 | 4.74 | 4.4 | 0.56 | 0.54 | 3.6 |
| **Misago** | 0.22 | 0.19 | 14 | 0.20 | 0.17 | 15 | 0.62/0.60 | 0.58/0.63 | 6.5/5.0 | 0.07 | 0.06 | 14 |
| **Zilver** | 1.46 | 1.59 | 8.9 | 0.76 | 0.73 | 3.9 | 0.90 | 0.93 | 3.3 | 0.30 | 0.31 | 3.3 |
| **Complete SE** | 1.14 | 1.28 | 12 | 1.24 | 1.36 | 9.6 | 1.66 | 1.68 | 1.2 | 0.28 | 0.29 | 3.6 |
| **EverFlex** | 1.06 | 0.93 | 12 | 1.13 | 1.30 | 15 | 2.24 | 2.31 | 3.1 | 0.36 | 0.41 | 14 |
| **Innova** | 0.95 | 1.04 | 9.5 | 0.73 | 0.62 | 15 | 1.26 | 1.44 | 14 | 0.26 | 0.29 | 12 |
| **Pulsar-18** | 0.34 | 0.29 | 15 | 0.28 | 0.25 | 11 | 0.50/0.050 | 0.52/0.06 | 4.0/17 | 0.07 | 0.06 | 14 |
| **LifeStent** | 0.56 | 0.59 | 5.4 | 0.56 | 0.65 | 16 | 1.79/1.76 | 1.73/1.91 | 3.4/8.5 | 0.18 | 0.22 | 22 |
| **S.M.A.R.T. Flex** | 8.70 | 9.89 | 14 | 1.06 | 1.05 | 0.9 | 3.54/3.50 | 3.52/3.13 | 0.5/11 | 0.37 | 0.44 | 19 |
| **Supera** | 0.28 | 0.23 | 18 | 0.22 | 0.20 | 9.1 | 6.40 | 2.54 | 60 | 0.21 | 0.22 | 4.5 |

**Figure 3.**
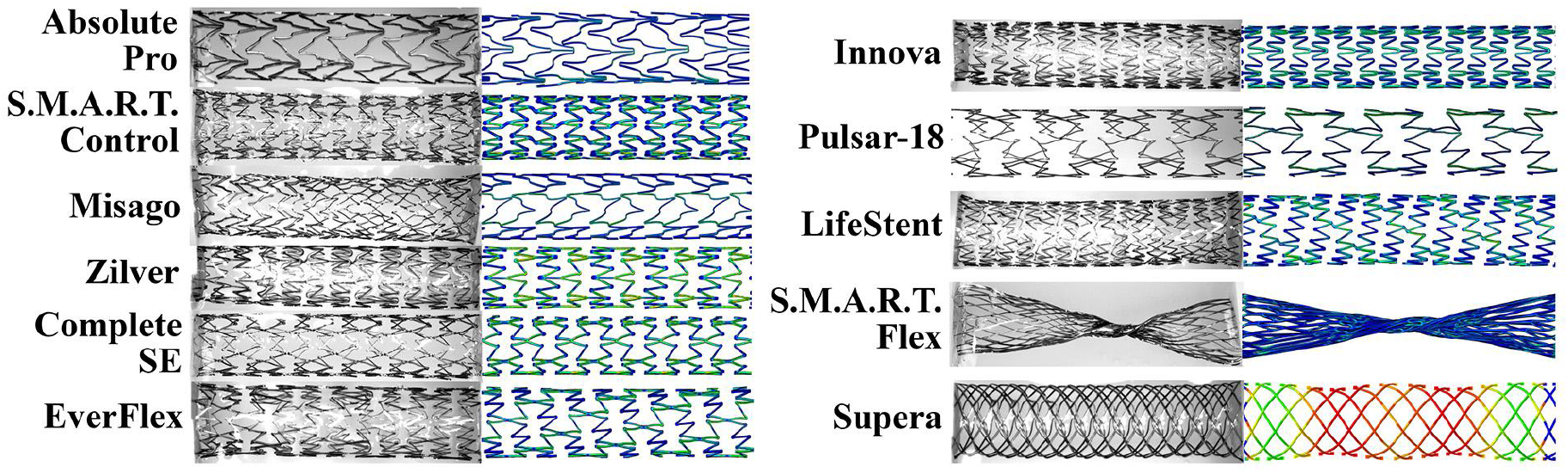
Experimental and FEA results for stents under the 25% axial tension load.

**Figure 4.**
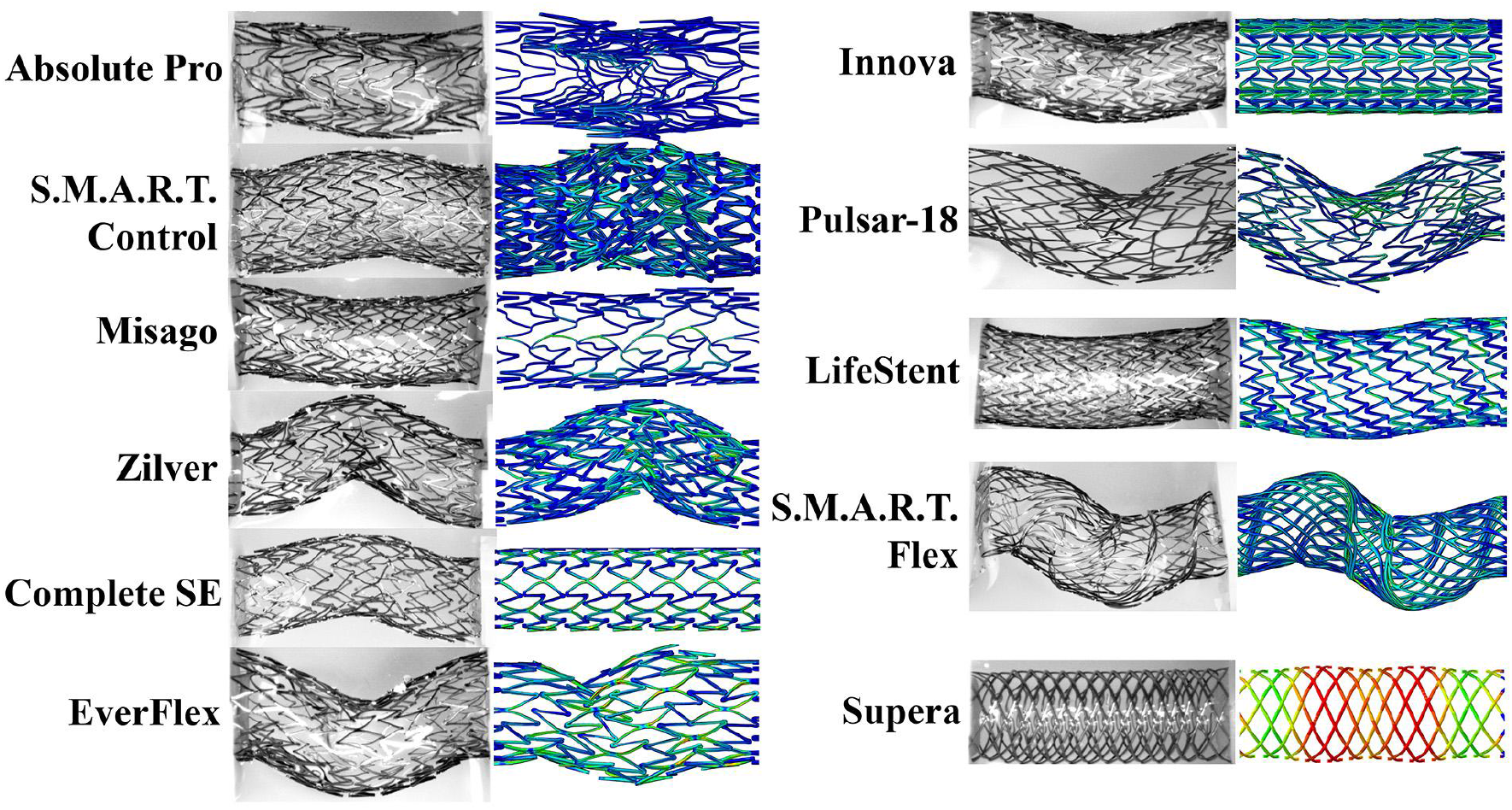
Experimental and FEA results for stents under the 25% axial compression load.

Under axial compression, S.M.A.R.T. Control demonstrated the highest reaction force (2.02 N), followed by Complete SE (1.24 N), while the lowest was again observed for the Supera (0.22 N) and Misago (0.20 N) devices. The FEA results were similar except for the LifeStent that demonstrated a slightly stiffer response than the Absolute Pro (0.52 N) and Innova (0.62 N) computationally, but not experimentally. The average difference between the FEA and the axial compression experiment was 10.1±4.9%, with the largest discrepancy observed for the LifeStent (relative error 16%, absolute error 0.09 N).

Figure 5 compares experimental and FE analysis of stents twisted 45°/cm and 90°/cm. Of all devices, Supera demonstrated the highest reaction moment (6.40 N•mm), which was 41% higher than the next stent S.M.A.R.T. Control (4.54 N•mm). The least resistance to twisting was observed for the Absolute Pro (0.54 N•mm) and Pulsar-18 (0.50 N•mm both clockwise and counter-clockwise) devices. Computational results followed the same trend, with the exception of Supera, for which the reaction force was underestimated (2.54 vs 6.40 N•mm), and Pulsar-18 stent that demonstrated a slightly larger reaction moment than the Absolute Pro computationally but not experimentally.

**Figure 5.**
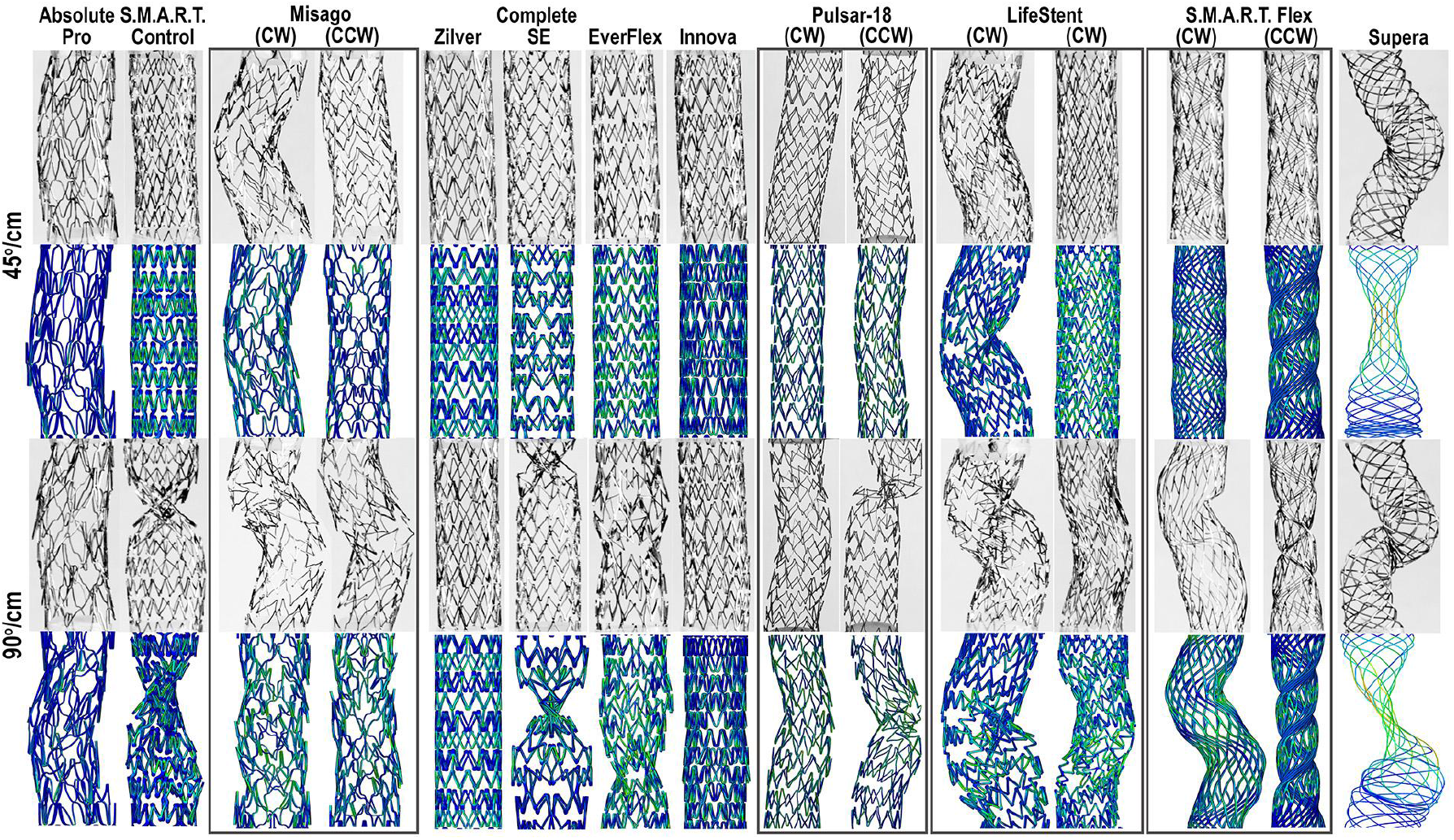
Experimental and FEA results for stents subjected to 45°/cm and 90°/cm twist. Clockwise (CW) and counter-clockwise (CCW) rotations are presented for non-symmetric devices.

The three-point bending comparison of the experimental and computational results is presented in Figure 6. Experimentally, the S.M.A.R.T. Control (0.56 N) demonstrated the largest reaction force, followed by the S.M.A.R.T. Flex (0.37 N) stent, while the lowest resistance to bending was demonstrated by the Pulsar-18 (0.07 N) and Misago (0.07 N) devices. FEA revealed the same trends, with an average difference of 10.6±6.6% across all devices, and the largest discrepancy for the LifeStent (22%), although the reaction force for this device was low both experimentally (0.18 N) and computationally (0.22 N). On average and across all deformations, the Supera (23±25%) and the LifeStent (13±8%) demonstrated the most difference between the experimental and the computational results, while the Zilver (4.9±2.7%) and the S.M.A.R.T. Control (5.2±2.1%) devices demonstrated the least.

**Figure 6.**
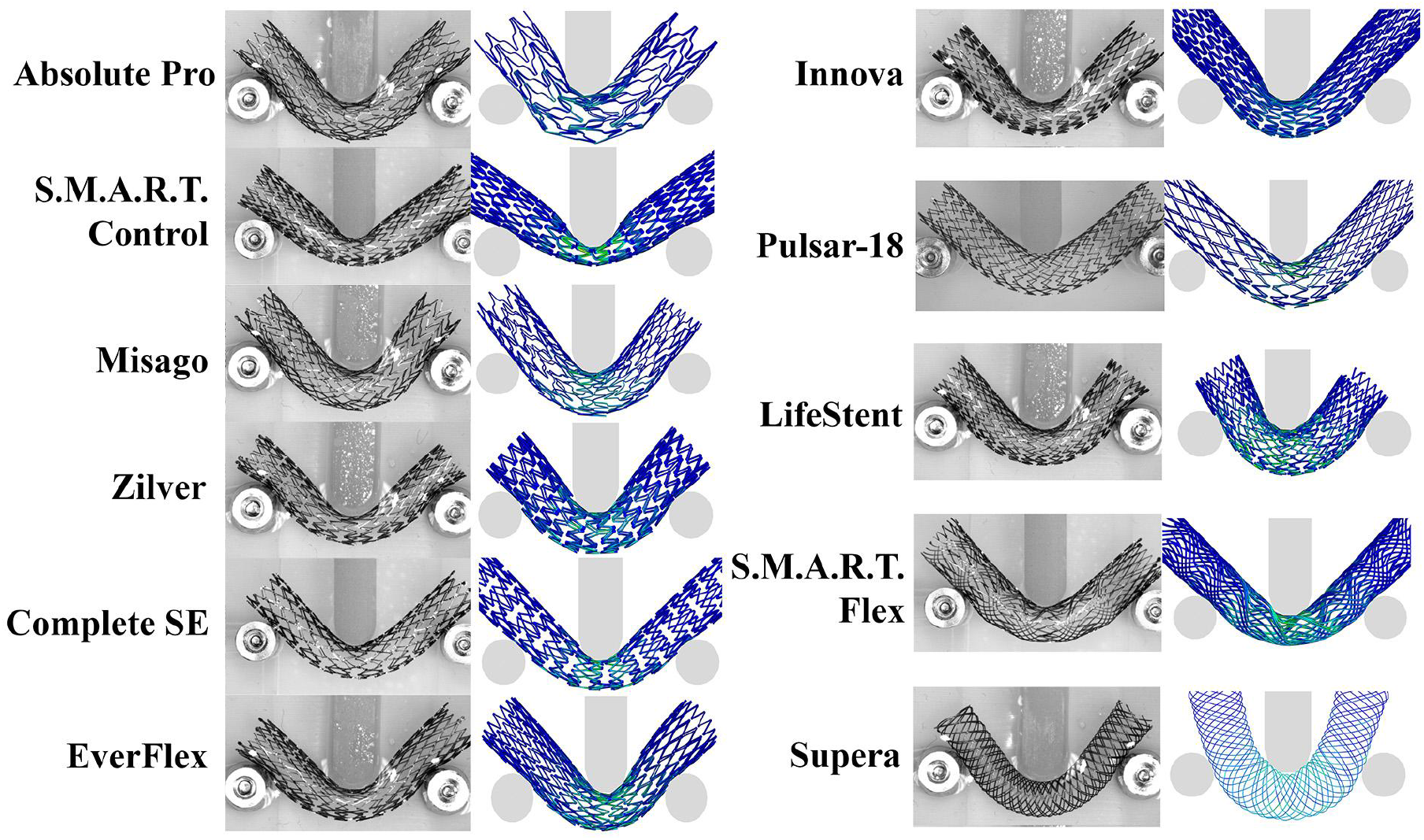
Experimental and FEA comparison of stents under the 3-point bending deformation.

Nitinol material parameters for all stents are presented in Table 3, and the stress-stretch curves generated using these parameters are plotted in Figure 7. Austenite Young’s moduli varied significantly across devices and ranged from 7.5 GPa for the S.M.A.R.T. Flex to 8.5 GPa for the Pulsar-18. The average austenite Young’s modulus across all stents was 32.5±24.1 GPa, while the average martensite Young’s modulus was 23.5±9.8 GPa. Transformation stretch ranged between 1.03 for the S.M.A.R.T. Flex and 1.08 for the Zilver. Figure 7 demonstrates that the material of the Zilver stent demonstrated the most compliant response, while Nitinol of the Pulsar-18 was the stiffest. Nitinol in the Cordis S.M.A.R.T. Flex had a significantly smaller hysteresis than in other stents, which likely stems from its significantly lower austenite Young’s modulus (7.5 GPa) compared with other devices. Material of the S.M.A.R.T. Control stent also demonstrated a small hysteresis, and its austenite Young’s modulus of 10 GPa was the second smallest in the cohort.

**Table 3.** Nitinol material properties for 11 peripheral stents determined using inverse computational analysis.

| Device | $E_A$, GPa | $E_M$, GPa | $\lambda^L$ | $t_L^S$, MPa | $t_L^E$, MPa | $t_{CL}^S$, MPa |
| --- | --- | --- | --- | --- | --- | --- |
| **Absolute Pro** | 20.0 | 47.8 | 1.063 | 432 | 451 | 900 |
| **S.M.A.R.T. Control** | 10.0 | 23.5 | 1.040 | 465 | 535 | 485 |
| **Misago** | 65.0 | 23.5 | 1.046 | 465 | 535 | 500 |
| **Zilver** | 12.5 | 15.0 | 1.082 | 386 | 411 | 150 |
| **Complete SE** | 35.0 | 23.5 | 1.046 | 435 | 505 | 582 |
| **EverFlex** | 40.0 | 30.0 | 1.046 | 465 | 535 | 582 |
| **Innova** | 20.0 | 15.0 | 1.040 | 465 | 535 | 500 |
| **Pulsar-18** | 85.0 | 23.5 | 1.060 | 465 | 535 | 580 |
| **LifeStent** | 37.5 | 23.5 | 1.050 | 375 | 425 | 582 |
| **S.M.A.R.T. Flex** | 7.50 | 10.0 | 1.030 | 465 | 535 | 500 |
| **Supera** | 25.0 | 23.5 | 1.046 | 465 | 535 | 582 |

**Figure 7.**
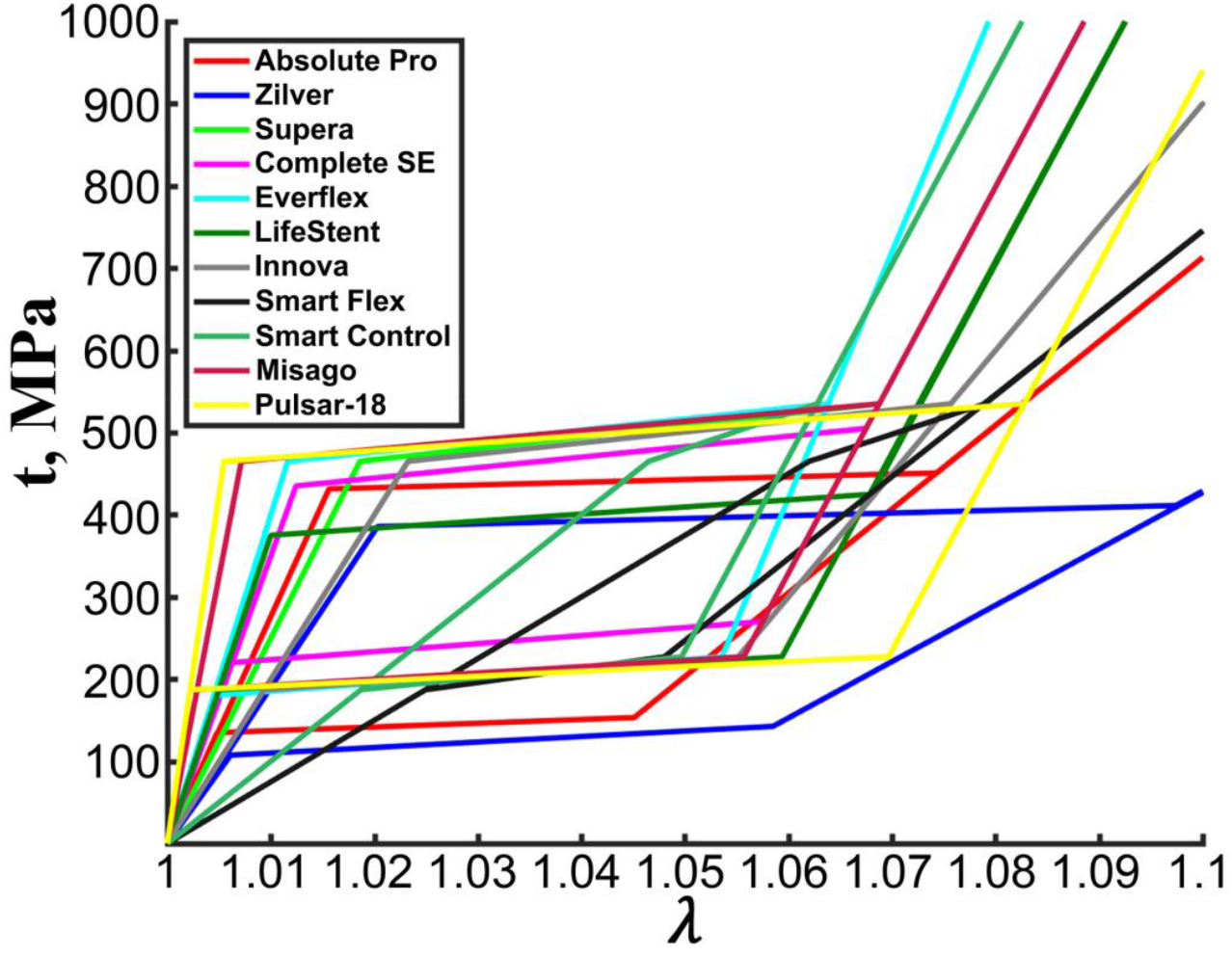
Nitinol stress-stretch plots for 11 stents determined using inverse computational analysis. Nitinol parameters used to generate the plots are summarized in Table 3.

Lastly, the stress-stretch curves obtained with the uniaxial extension of the straight connector bridge struts from the Absolute Pro and Zilver stents, overlaid with their computationally-determined counterparts, are demonstrated in Figure 8. The experimental curves are in good agreement with the FEA-determined values and illustrate a typical superelastic response of the Nitinol material. Experimentally, the Absolute Pro demonstrated significantly higher austenite (nearly two-fold) and martensite (more than threefold) Young’s moduli compared to the Zilver, as well as a higher start of transformation stress in loading (482 MPa vs 386 MPa). Plateau regions from the transition from austenite to martensite were similar for the two devices, although somewhat longer for the Zilver and reaching nearly 1.10 stretch, while the Absolute Pro began its martensitic phase at 1.07. At 1.10 stretch, the Absolute Pro demonstrated nearly two-fold greater stress than Zilver (786 vs. 462 MPa), but the Zilver stent demonstrated a 23% higher transformation stretch than the Absolute Pro (1.08 vs. 1.06). In terms of unloading, Zilver began the transition from martensite back to austenite sooner (1.10 vs. 1.07), but the plateau region in unloading appears similar in terms of stress (164 MPa for the Absolute Pro vs. 147 MPa for the Zilver). Nevertheless, Zilver’s transition plateau region ended at 1.03 stretch, while the Absolute Pro’s recovered to 1.01. Notably, Zilver did not fully unload to a stretch of 1.0, which was likely a result of minor slippage of the strut from the epoxy base. The Absolute Pro demonstrated a start and end of transformation stress during unloading of 153 MPa and 135 MPa, while the Zilver demonstrated values of 143 MPa and 108 MPa, respectively.

**Figure 8.**
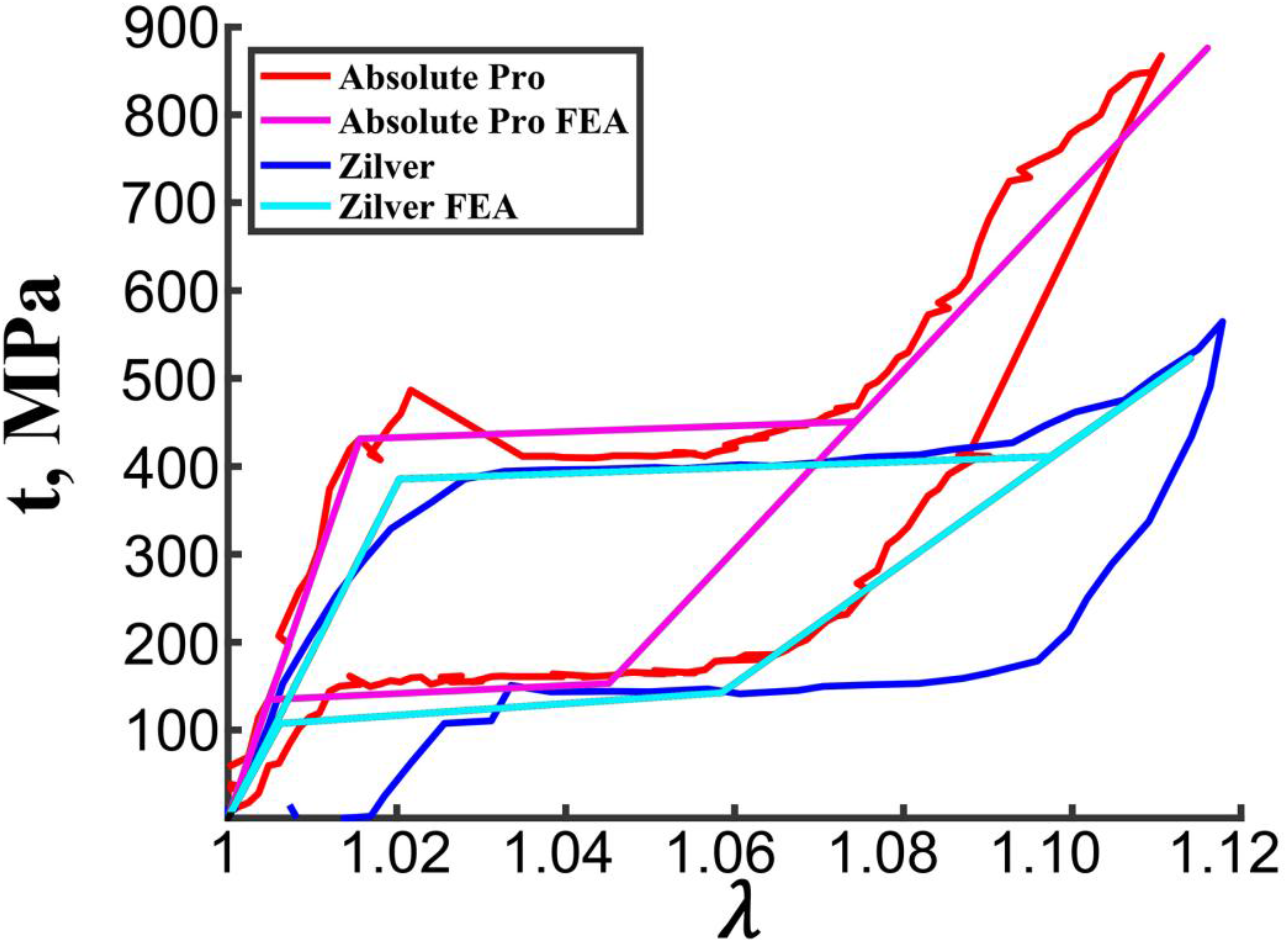
Experimental stress-stretch curves from the uniaxial extension test of the straight connector bridge struts from the Absolute Pro (red) and Zilver (blue) stents overlaid with their respective FEA-determined curves (magenta and cyan).

## 4. DISCUSSION

Stent-artery interactions resulting from limb flexion-induced deformations^13–15,37–39^ are thought to contribute to poor clinical outcomes of PAD repair^22,40–42^ and the high rate of stent fractures^12,43–46^. Computational modeling can help identify stent design features that result in better device biomechanics and hemodynamics^47^, but to produce realistic results, these models need accurate inputs on device mechanical properties. Most peripheral stents are made of superelastic Nitinol material^21^, which is a shape memory alloy with unique characteristics that allow the stent to expand to a pre-set shape once released from the catheter without the assistance of a balloon, and return to this shape after being deformed during limb flexion. The mechanical characteristics of Nitinol also allow “biased stiffness”, which means that a stent that recovers from the crimped position is more resistant to compression than to expansion, and since stents are usually oversized to account for the overstretched lumen after angioplasty, “biased stiffness” allows reducing the outward force. Different manufacturing and postprocessing techniques^21,33,48^, such as heat treatment temperature and time, cooling rate, or electropolishing, are used to influence the microstructure of Nitinol and achieve different mechanical characteristics. But since manufacturers seldom disclose their Nitinol processing methods or material properties^32^, many studies use “generic” Nitinol parameters^23–34^ in their simulations, which often has a significant effect on the results. Kleinstreuer et al.^30^ conducted a computational study comparing two different sets of Nitinol material parameters used for tubular diamond-shaped stent-grafts under cyclic loading, and reported markedly different behavior. Similarly, another study^31^ compared finite element simulations using various Nitinol parameters from the literature and demonstrated that material inputs significantly affected stress-stretch calculations. These and other similar studies highlight the importance of having correct material parameters for computational models, but their direct experimental assessment remains limited.

Several studies have used uniaxial tensile testing of straight wires^27,35,49^ or tubes^28^ to assess Nitinol behavior, but these approaches are largely not applicable to peripheral stents that often have complex unit cell geometries with an exceedingly limited number of straight segments that can be mechanically tested. Of the 11 stents evaluated in our study, only two devices had suitably long straight sections that could be isolated and studied under uniaxial tension. These devices were Abbott Absolute Pro and Cook Zilver stents, and their stress-stretch responses were markedly different, supporting the importance of determining device-specific material parameters. Using these two tests as a benchmark, we have assessed Nitinol characteristics of other stents with more complex geometries using inverse computational analysis coupled with benchtop evaluation of these devices under axial tension, compression, bending, and torsion deformations.

Inverse computational analysis is instrumental in a variety of problems where direct experimental measurement is difficult. For example, it is used to determine the manufacturing shape of wind turbines based on service loads^50^, assess the material properties of the aortic wall^51^ or ovine mitral valve anterior leaflet from the *in vivo* measured displacements^52^, and is widely implemented in the aerospace industry for real-time control and health monitoring of structures based on surface-measured strains^53^. The major challenge of using the inverse analysis is that boundary conditions are often poorly defined, and the material properties are complex and are described by multiple variables that need to be determined from limited experimental data^54^. To mitigate these issues, we have performed controlled benchtop experiments that included all major modes of PAD stent deformation and have validated our results with single-strut testing of two stents.

Our data demonstrate that Nitinol material properties are widely different across devices but generally fall in the range reported in the literature. Specifically, we found an average Young’s modulus of 7.5-85 GPa (average 32.5±24.1 GPa, austenite) and 10-47.8 GPa (average 23.5±9.8 GPa, martensite), while Young’s modulus reported in the literature ranges anywhere from 30 to 68 GPa (austenite) and from 18.5 to 51 GPa (martensite). Nevertheless, several stents in our study demonstrated an austenite Young’s modulus values below 30 GPa, including the Absolute Pro which was evaluated using the single strut experiment. In terms of the volumetric stretch, our values ranged 1.03-1.08, while values reported in the literature were 1.04-1.06. The Cook Zilver stent, which was also evaluated using the single strut experiment, demonstrated the largest volumetric stretch (1.08), exceeding literature values.

When comparing stents made by the same manufacturer, the S.M.A.R.T. Control and the S.M.A.R.T. Flex (Cordis) devices showed similar results with high reaction forces and moments and low austenite Young’s moduli. The Complete SE and the EverFlex (Medtronic) stents also demonstrated similar responses in terms of reaction forces and moments, and had similar Nitinol material properties. Lastly, the Absolute Pro and the Supera (Abbott Vascular) demonstrated similar reaction forces for all deformations, but the Supera had a nearly 12-fold higher reaction moment under experimental torsion than the Absolute Pro. Their Nitinol characteristics were also different. While dissimilarity in device behavior clearly stems from design (i.e., Supera is braided and Absolute Pro is laser-cut), the differences in Nitinol properties may in part be due to the fact that Supera was not originally developed by Abbott but was initially manufactured by IDEV (IDEV Technologies Inc, Webster, TX, USA).

The inverse FEA method used in our study was effective in determining Nitinol material properties for commercial peripheral stents, and the summarized parameters can inform computational models of device-artery interactions. Nevertheless, these results should be viewed in the context of study limitations. First, reverse-engineered stent geometry may be imprecise, which would translate into differences in the determined mechanical characteristics. For this reason, we have not attempted to determine Nitinol parameters with precision greater than the 15% combined relative average error in all four deformation modes between the FEA and the experiment, particularly since many devices that demonstrated appreciable relative error had a small absolute error. Second, we have only tested one device of each kind. Though we do not expect the variation between the same devices to be significant, studies including a larger number of samples would allow a more accurate determination of Nitinol parameters, in part because setting up the experiment and attaching the stent to the testing apparatus can also introduce variations. Third, while we were able to determine the main Nitinol parameters that primarily define stent behavior, they may not necessarily be unique, and there could exist other combinations of these parameters that would produce similar results. While these and other limitations of this work are being addressed, we hope that our study would be useful for computational simulations of peripheral stent-artery interaction, which may help improve the outcomes of endovascular PAD repair.

## ACKNOWLEDGEMENTS

This work was supported in part by the National Institutes of Health under award numbers HL125736, HL147128, and P20GM152301.

## Notes

### Competing Interest Statement

The authors have declared no competing interest.

